# Integrated Spatial Analysis of Ovarian Precancerous Lesions

**DOI:** 10.1101/2025.06.24.661327

**Authors:** Tu-Yung Chang, Yen-Wei Chien, Szu-Hua Chen, Annelise Sokolow, Yeh Wang, Brant G. Wang, Tricia Numan, M. Herman Chui, Rebecca Stone, Thomas Pisanic, Nicholas Papadopoulos, Tian-Li Wang, Christopher Douville, Leslie Cope, Ie-Ming Shih

## Abstract

Studying precancerous lesions is essential for improving early detection and prevention, particularly in aggressive cancers such as ovarian carcinoma. Here, we conducted integrated and spatial analyses of transcriptomes, aneuploidy, and clinic-pathological features in 166 ovarian precancerous lesions. Four pre-cancerous subtypes were identified transcriptomically: proliferative, immunoreactive, dormant, and mixed. These subtypes varied in their frequency of germline-*BRCA1/2* mutations, aneuploidy, *CCNE1*/*MYC* amplification, proliferative activity, immune-regulatory gene expression, and histological features. Notably, the immunoreactive subtype upregulated immune-regulatory genes, exhibited chronic inflammation, and was enriched in cases with germline-*BRCA1/2* mutations, deletions of chromosomes 17 (harboring *TP53* and *BRCA1)* and 13 (harboring *BRCA2*), leading to a double “two-hit” involving *TP53* and *BRCA1/2*. Tumor invasion was associated with the activation of interferon response pathways, epithelial-mesenchymal transition, and extracellular matrix remodeling. In summary, our results elucidate the earliest molecular landscape of ovarian precancerous lesions, serving as the foundation for future risk stratification to identify aggressive pre-cancerous lesions.

## Introduction

Most solid tumors evolve from their pre-cancerous stages, where multiple clonally developed precancerous lesions arise. These pre-cancerous lesions may regress, remain stable, or progress. Elucidating their pathogenesis provides novel insights into the primordial niche of tumor initiation and reveals the molecular mechanisms that drive tumor progression. Such knowledge is essential for developing clinical strategies to intercept cancer at earlier, more treatable stages, which is particularly crucial for aggressive cancer types that often evade early detection, such as ovarian carcinomas.

One of the “hard problems” in eradicating ovarian cancer is that most patients with high-grade serous carcinoma (HGSC), the most common type of epithelial ovarian cancer, are diagnosed at advanced stages. Current treatment interventions are largely ineffective due to recurrence, making cure an elusive goal despite recent improvements in cytoreductive surgery, chemotherapy, and PARP1 inhibitor maintenance therapy.

Historically, HGSC has been regarded as arising *de novo* from the ovaries; thus, the canonical term “ovarian” serous carcinoma has been used for decades. A “crisis of confidence” in this assumption has emerged due to the limited evidence demonstrating that HGSC precursors actually originate from the ovarian surface epithelium ^1^. More than twenty years ago, a study performed meticulous microscopic examination of adnexal tissues from women with inherited *BRCA1* pathogenic mutations and reported the presence of dysplastic epithelial cells, indicative of HGSC precursor lesions, in the fallopian tubes, rather than in the ovaries themselves ^2–4^. In diagnostic pathology, these HGSC precursor lesions were later termed serous tubal intraepithelial carcinoma (STIC) and serous tubal intraepithelial lesion (STIL)-a variant which does not meet all morphologic or immunohistochemical criteria diagnosis of STIC ^5^.

Subsequently, several research teams have provided evidence to independently support the notion that most “ovarian” HGSCs originate from microscopic precursor lesions of the fallopian tubes ^6–8^. Population-based studies indicate that salpingectomy alone significantly reduces the incidence of ovarian cancer ^9–12^. In addition to clinicopathological evidence, multiple studies have demonstrated that the fallopian tubes are likely the tissue of origin for most HGSCs. Transcriptomic analysis indicates that most HGSCs have a closer molecular relationship to fallopian tube epithelium than to ovarian surface mesothelium ^13^, and mutational analysis has identified a related clonal relationship between STICs and concurrent HGSCs ^14–16^.

The paradigm shift regarding the origin of HGSC from the ovaries to the fallopian tubes raises several critical research questions: What are the key molecular alterations driving tumor initiation and progression to HGSCs? How do genetic predisposition (e.g., *BRCA1/2* germline mutations), molecular phenotypes, morphological features, and the immune landscape interact in HGSC precursor lesions? Are HGSC precancerous lesions a heterogeneous group, and if so, how can we classify them into biologically distinct subtypes? Can we identify a precancerous subgroup that is responsible for disseminated HGSC years after risk-reducing bilateral salpingectomy? In fact, risk stratification has become a critically unmet need in the clinical management of women diagnosed with STICs because they appear to face an unresolved threat of developing HGSC of the peritoneum years later and effective mitigation strategies remain undefined.

Pilot spatial analyses have reported alterations in genome-wide DNA copy numbers, DNA methylation, and whole transcriptomics, in addition to mutations identified by whole-exome sequencing in HGSC precancerous lesions ^17^. However, an integrated analysis has not been performed to correlate various molecular and clinicopathological data, which would yield much more holistic, clinically actionable understanding of biology in ovarian cancer development. This study represents a concerted effort to tackle these essential questions through an integrated spatial analysis of transcriptomics, aneuploidy patterns, morphological features, and clinical information in a relatively large cohort of specimens consisting of 438 spatial regions of interest, including 166 discrete precursor lesions. Importantly, we provide our data along with a user-friendly portal to facilitate public access and data inquiry.

## Materials and Methods

### Tissue collection and preparation

Fallopian tube tissues diagnosed with STICs, STILs, and p53 signatures were retrieved from the pathology archive at Johns Hopkins Medical Institutions (Baltimore, MD), Inova Fairfax Hospital (Fairfax, VA), and Memorial Sloan Kettering Comprehensive Cancer Center (New York, NY) between 2018 and 2023. For laser capture microdissection (LCM), slides were cut to a thickness of 10 μm, while slides for the GeoMx Digital Spatial Profiler platform were cut to a thickness of fourLμm. The study was conducted in accordance with the ethical principles outlined in the Declaration of Helsinki. It was approved by the Institutional Review Boards (IRBs) of Johns Hopkins Medical Institutions (IRB No. 00147713 and 00052121), Inova Fairfax Hospital (IRB No. U20-07-4179), and Memorial Sloan Kettering Cancer Center (IRB No. 16-1684).

### Histologic Review of Diagnosis and Laser Capture Microdissection

A total of 68 patients with 294 precancerous epithelial samples consisting of 162 STIC/STIL and four p53 signatures, along with 29 concurrent carcinoma lesions, 99 paired histologically unremarkable or normal fallopian tube epithelia (NFT), and 144 diagnosis-paired stromal samples, were included in this study. Two morphological subgroups were defined based on criteria from a recent report: ^18^ a BLAD subgroup characterized by budding, loosely adherent, or detached cells in over 10% of the lesion, and a Flat subgroup, with more than 90% of the lesion lacking these features. All tissue sections were scanned into a digital pathology platform using a Hamamatsu-S60 slide scanner. Stromal lymphocyte density (enriched versus non-enriched) was graded by two pathologists (TC and IS) who reached a consensus. A lymphocyte enriched lesion was defined as a lesion with significantly higher lymphocyte density in the lesional stroma than in the background fallopian tube mucosa from the same tissue sections.

We performed laser capture microdissection (LCM) on each area of interest. The slides were deparaffinized using xylene prior to LCM. Each selected region contained a single, continuous lesion; spatially separated lesions were considered distinct samples. DNA was purified with the Qiagen QIAamp DNA FFPE Advanced Kit, and DNA concentration was quantified using a Qubit fluorometer.

### Spatial Transcriptomic Profiling

To measure mRNA expression levels in paraffin-embedded tissues, we employed the GeoMx Human Whole Transcriptome Assay (Nanostring Technologies, Inc.). A total of 442 areas of interest (AOIs) were selected from individual digitized fluorescent images based on morphological features and ancillary markers, including pan-cytokeratin, p53, and DAPI, under the supervision of a pathologist. The methods, including bioinformatics analysis of Nanostring GeoMx, were detailed elsewhere ^19^. Following AOI selection, ultraviolet (UV) light was applied to each AOI to release RNA identification-containing oligonucleotide tags from the CTA probes, which were then collected for sequencing. Among the 442 AOIs, we included 49 AOIs from our previously reported study ^17^, comprising epithelial and stromal components.

### Batch Correction and Non-negative Matrix Factorization

All data analyses were performed using R (version 4.4.1 , RRID:SCR_001905) and RStudio (version 2023.12.1 , RRID:SCR_000432). Raw count data were initially corrected for batch effects using the remove batch effect function from the limma R package ^20^, treating each GeoMx run (corresponding to four slides) as a separate batch, followed by a variance-stabilizing transformation. Batch correction was assessed using kBET scores ^21^ and PCA visualization. To classify epithelial samples, we applied Non-negative Matrix Factorization (NMF) with the NMF R package ^22^, performing 100 consensus runs to ensure clustering stability. The optimal number of clusters was selected based on both quantitative measures of clustering quality and biological interpretability, including metrics such as cophenetic correlation, dispersion, and silhouette width.

### Differential Expression and Gene Set Enrichment Analysis

The limma R package was used to identify differentially expressed genes (DEG) across the four proposed molecular subtypes. To explore the biological significance of the identified DE genes, we applied the Gene Set Enrichment Analysis (GSEA) ^23,24^ using curated DEG sets.

### Repetitive Element Aneuploidy Sequencing System (RealSeqS)

We applied RealSeqS to assess aneuploidy using the DNA (2.5 ng) extracted from the laser capture microdissected tissues. The method of RealSeqS has been detailed in a previous report^25^. used a single primer pair purchased from IDT, Inc. (Coralville, Iowa) to amplify approximately 350,000 genomic loci. After the initial round of PCR, a second round was performed to incorporate dual indexes (barcodes) into each PCR product, which was purified with AMPure XP beads (Beckman cat # a63880) before sequencing and the exclusion of fragment sizes <100 bps. Sequencing was performed on an Illumina HiSeq 4000 using single-end reads of 150 bps length.

The method used in this study to process the raw sequencing data is available at https://zenodo.org/record/3656943#.YaZZCdDMKUk. A complete list of the repetitive elements as well as the bioinformatic pipeline downloadable link were available in Dataset S01 from ^25^. The bioinformatics pipeline generates a file with the observed counts at each locus. The following Python2 dependencies are required for the bioinformatics pipeline: argparse, json, logging, multiprocessing, and re. We used the following versions when processing the raw data: Python 2.7.14, argparse 1.1, json 2.0.9, logging (0.5.1.2), multiprocessing (0.70a1), and re (2.2.1).

Chromosomal alterations are identified when a sample’s normalized read counts within a genomic region significantly differ from what is expected in an euploid sample. Each sample is compared to a reference panel consisting of 30 euploid genomic DNA samples from individuals without cancer. These samples were epithelial cells collected and purified from the voided urine of young individuals (aged <25 years) and were not included in the study. RealSeqS was performed on those 30 samples before evaluating the experimental samples in our study. Each experimental sample was matched to a smaller subset (n=7) of the larger reference panel (n=30) that was most similar in terms of the amplicon distributions generated by RealSeqS based on Euclidean distance. The statistical significance for the 39 non-acrocentric chromosome arms was then calculated using the equation below.

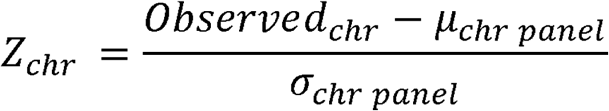

An arm was classified as aneuploid if |Z_chr_|>5, gained if Z_chr_ > 5, and lost if Z_chr_ < -5. In addition to generating arm-level statistical significance, the circular binary algorithm (CBS) as distributed in the DNA copy library (v1.58) within the R (v3.6.1) programming language was applied to 50-kb non-overlapping genomic intervals for each experimental sample ^26^. The CBS algorithm identifies sub-chromosomal focal alterations. A focal amplification for 19q12, 19q13.2, or 8q24 was called if CBS produced a sub-chromosomal segment surrounding the region of interest and had a log2 ratio > 0.25. We successfully conducted Real-SeqS on 51 samples containing sufficient DNA amounts after laser capture microdissection of the targeted lesions. The current study combined 31 new DNA samples and 20 previously reported RealSeqS data for analysis ^27^.

### Aneuploidy Index and Molecular Distance Calculation

The aneuploidy index was calculated by summing the number of base pairs with a log_2_ ratio greater than 0.25 (indicating copy number gain) or less than -0.25 (indicating copy number loss) across the genome. We assessed the molecular similarity between samples using molecular distance, which was calculated with the Euclidean distance metric based on gene expression profiles. Gene counts were first filtered to include the top 50 uniquely upregulated genes from each molecular subtype. For reference, gene expression profiles from histologically unremarkable tubal epithelium, ovarian mesothelium, and HGSC samples were included in the distance calculations.

### Integration of RNA and DNA Data via Cytoband Mapping

We investigated the relationship between gene expression and genomic alterations by integrating RNA and DNA data according to chromosomal cytoband regions. Genes from the RNA-seq data were annotated with their corresponding cytoband locations, and expression patterns were compared to DNA copy number alterations within the same areas derived from sequencing data. Pearson correlation was calculated for each gene between its expression levels and corresponding DNA segment log_2_ ratio values across matched samples. This analysis assessed the relationship between regional transcriptional changes and underlying genomic gains or losses, enabling the identification of cytobands with concordant alterations at both the RNA and DNA levels.

### Immunohistochemistry

Immunohistochemistry (IHC) was performed to evaluate protein expression levels. Five slides from the original cases, along with 10 additional cases using two HGSC tissue microarray slides, were used. Immunohistochemistry was carried out using the previously described methods 28,29. Briefly, formalin-fixed, paraffin-embedded tissue sections were first deparaffinized and rehydrated. Antigen retrieval was conducted using the Trilogy solution (Cell Marque, #920P-10). All tissue sections were quenched with 3% H₂O₂ to inactivate endogenous peroxidase activity. Sections were incubated with primary antibodies at 4°C overnight.

The following antibodies were used for verification: anti-SOX4 monoclonal antibody [Abcam, CL5634, RRID:AB_3083741], anti-MCM7/PRL monoclonal antibody [Abcam, EP1974Y, RRID:AB_881187], anti-KIFC1 monoclonal antibody [Abcam, 11445, RRID:AB_2827938], and anti-DBN1 polyclonal antibody [Sigma, HPA051452, RRID:AB_2681489]. In all cases, either the original stained slides or additional immunostaining using p53 [Roche, Bp53-11] and Ki67 [Roche, 30-9] was utilized. Visualization of immunoreactivity was achieved using the DAKO EnVision+ System-HRP with goat anti-rabbit or anti-mouse IgG and the DAKO DAB+ Substrate Chromogen System. We used open-source software QuPath 30 to quantify the percentage of Ki67-positive epithelial cells (the Ki67 index) and relative protein expression in immunohistochemistry in digitized images. This method (H-score) provided scores for individual tissue areas by calculating the sum of the stain intensity score (0-3) multiplied by the percentage of epithelial cells from each intensity category.

### Statistical analysis

Comparisons across molecular subtypes were conducted using chi-square tests or Fisher’s exact tests, depending on expected cell counts. Pearson correlation was used for association analysis between gene expression and copy number data, with Benjamini–Hochberg correction applied for multiple testing. Differential expression was evaluated using the limma package, with significance defined as an adjusted p-value < 0.05 and absolute log_2_ fold change > 0.5.

### Data Availability

The spatial transcriptomic data generated and analyzed in this study are publicly available in the Gene Expression Omnibus (GEO) under accession number GSE298031.

## Results

### Spatial Transcriptomic Profiles in Ovarian Cancer Precursor Lesions

We applied the GeoMx® DSP platform to analyze whole transcriptomes in 438 areas of interest (AOI), which included 294 epithelial AOI and 144 stromal AOI located immediately beneath specific epithelial AOI. These AOI represent a cohort of 68 women. Based on histopathological diagnoses, we curated 162 AOI from morphologically defined STICs (112 precursors without HGSC and 50 precursors with concurrent HGSC), four from p53 signatures, 29 from HGSCs that had concurrent STICs analyzed, and histologically unremarkable fallopian tube epithelium adjacent to the lesions. The study design is outlined in Fig. 1A.

**Figure 1.**
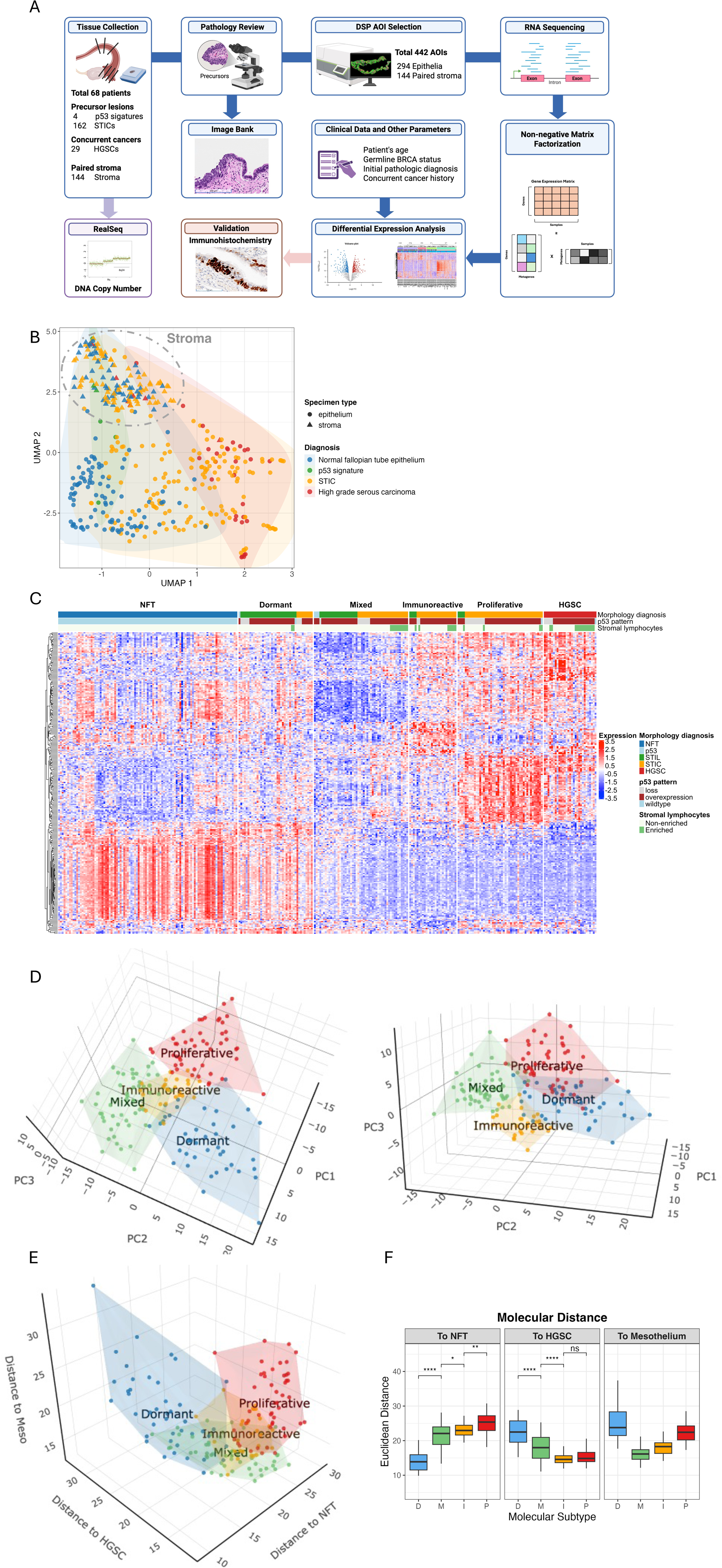
Spatial transcriptomic analysis identifies molecular heterogeneity of tubal pre-cancerous lesions. (A) A schematic illustration of the study design. (B) UMAP visualization of transcriptomics from epithelial and stromal samples of different diagnostic groups. (C) The heatmap displays hierarchical clustering of the top 50 differentially expressed genes in tubal pre-cancerous (STIC) lesions and cancerous lesions. Four STIC molecular subtypes are identified. (D) 3D PCA analysis demonstrate the separation of STIC subtypes (or pre-cancerous lesions of Fallopian tube). Plots are generated based on the leading principal components. (E) 3D plots illustrate the molecular distances of STIC subtypes and the referenced normal fallopian tube epithelium (NFT), high-grade serous carcinoma (HGSC), and ovarian mesothelium. (F) Boxplots present the molecular (Euclidean) distances between individual STIC subtype and the referenced NFT (left), HGSC (middle), and ovarian mesothelium (right).

First, we applied UMAP and a heatmap to assess molecular relationships in individual AOI across different tissues (Fig. 1B and Fig. S1). As anticipated, epithelial and stromal cells were transcriptomically distinct, demonstrating rigor in our AOI selections. The stromal AOI, regardless of their corresponding epithelial lesions, clustered tightly (Fig. 1B). Since unsupervised heatmaps did not detect distinct clusters in the stromal component as they did in the epithelial component among STICs (Fig. S1A and S1B), we compared the stromal transcriptomes of STICs as a whole to those of HGSCs. As shown in Fig. S2A, we identified 15 differentially expressed genes that were upregulated in HGSC stroma, primarily associated with tumor-associated M2-like macrophages. Additionally, stromal lymphocytes increased in HGSC stroma, and pathway analysis revealed activation of the interferon-γ and interferon-α response pathways (Fig. S2B, S2C).

As our preliminary results did not reveal substantial and/or consistent transcriptomic differences based on GeoMx® DSP analysis, we chose to focus our study on analyzing epithelial AOI. From these results, we observed that STIC lesions and HGSC samples were significantly distinct from their corresponding normal fallopian tube epithelium (Fig. 1B). STIC lesions were more widely dispersed than their stromal counterparts, suggesting the presence of transcriptomically-distinct STIC subgroups. This finding is of great interest because attempts to histologically classify STIC lesions into various subtypes have been challenging, with unsatisfactory reproducibility even among experienced pathologists ^31^. This highlights a critical need for a molecular classification system that can help refine the pathological diagnoses of STICs in clinically meaningful ways.

### Tubal Precancerous Lesions are Molecularly Heterogeneous

Based on the results shown in Fig. 1B and Fig. S1, we employed non-negative matrix factorization and identified four molecular subtypes of STIC. These subtypes displayed distinct gene expression profiles, as demonstrated by hierarchical clustering in the heatmap and principal component analysis (PCA) (Fig. 1C and 1D). We labeled these subtypes as proliferative, immunoreactive, mixed, and dormant STIC subtypes (hereafter referred to as PIMD). In addition to separating STICs in the gene expression-based PCA space, we further evaluated the PIMD subtypes in the cell-based PCA space, assessing their similarity to normal fallopian tube epithelium, ovarian surface mesothelium, and HGSC cells. Our results indicated that the PIMD subtypes were also well defined in this domain (Fig. 1E). A simplified box plot illustrated that the dormant STIC subtype and the proliferative subtype were molecularly closest and farthest from the normal fallopian tube epithelium, respectively. In contrast, the proliferative subtype and the dormant subtype were molecularly closest and farthest from HGSCs, respectively (Fig. 1F).

Volcano plots identified differentially expressed genes (DEGs) that characterized individual PIMD subtypes. Ranks of the pathway activation or inactivation were also presented according to DEGs. As shown in Fig. 2A, the proliferative STICs upregulated several tumor-promoting DEGs, including *NACC1, NOTCH3*, *IGFBP2*, *LAMC1*, *STMN1*, *SOX4*, *SOX17*, *BCAM, FASN*, and others frequently associated with HGSCs ^32–37^. Immunoreactive STICs more frequently upregulated various HLA immune regulatory genes, including *HLA*, *TAP2*, *LCN2*, *A2M*, *GPNMB,* and genes involved in the interferon response pathway (Fig. 2B). In contrast to the other STIC subtypes, the dormant subtype frequently upregulated fallopian tube differentiation markers, including the secretory marker, *OVGP1,* and ciliated cell differentiation markers, *FOXJ1* and *CAPS*. Upregulation of these markers has been reported to be associated with improved overall survival in ovarian HGSC patients (Fig. 2C) ^38,39^. Additionally, *OVGP1* and *CRISP3,* which were elevated in dormant STICs, have been shown to be downregulated in HGSC. Interestingly, the mixed subtype did not exhibit many upregulated DEGs compared to other subtypes; instead, these STICs displayed decreased expression levels of various genes relative to the different subtypes (Fig. 2D). Comparing individual PIMD subtypes with histologically unremarkable fallopian tube epithelium revealed pathway activations, as shown in Fig. S3A. By comparing dormant STICs to the normal tubal epithelium, we found upregulated DEGs involved in pathways related to Myc targets, E2F targets, and G2M checkpoint (Fig. S3B and S3C).

**Figure 2.**
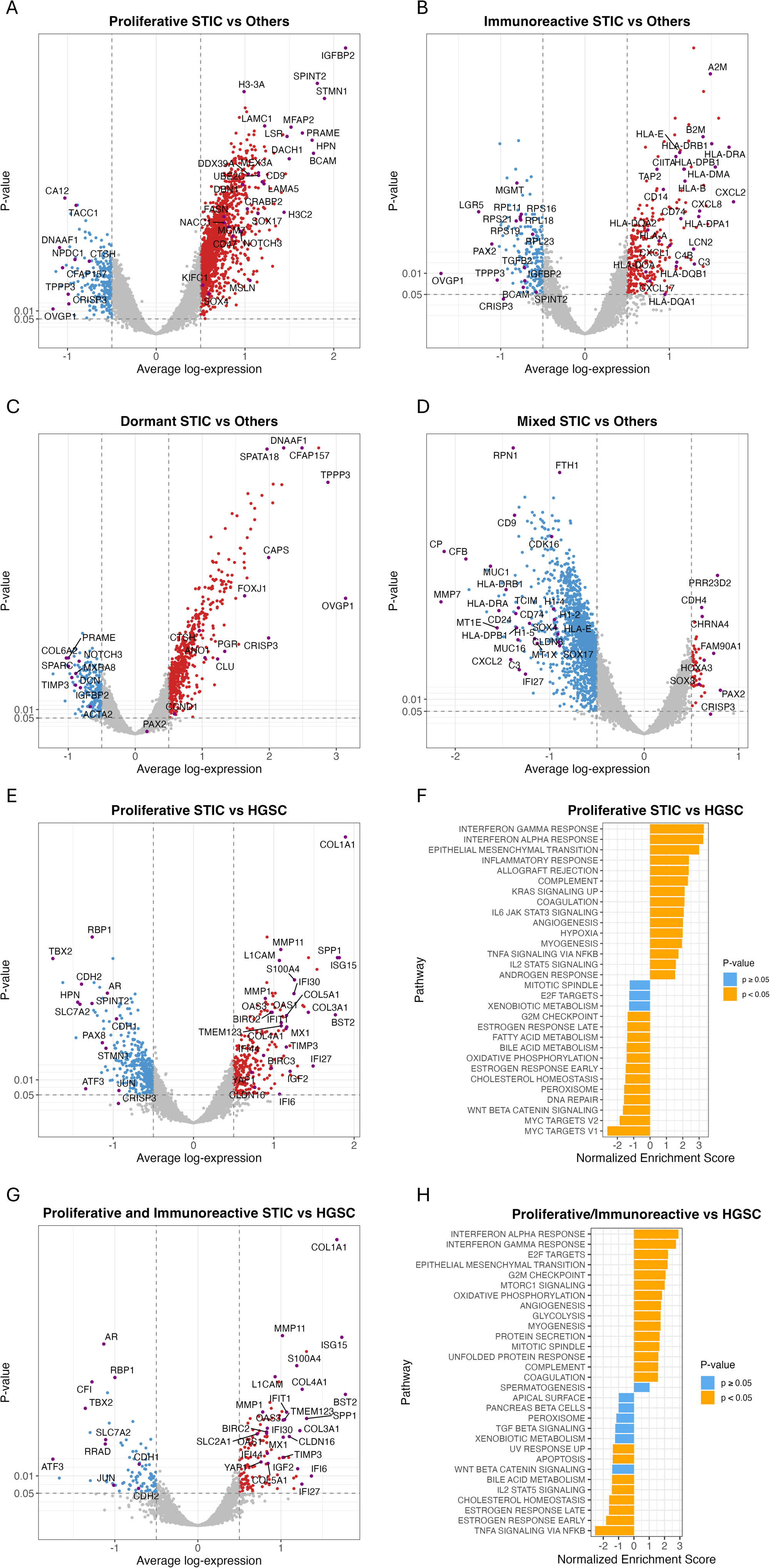
Differentially expressed genes across STIC molecular subtypes. (A) Differential gene expression in the Proliferative subtype. (B) Differential gene expression in the Immunoreactive subtype. (C) Differential gene expression in the Dormant subtype. (D) Differential gene expression in the Mixed subtype. (E) Differential gene expression in the Proliferative STIC subtype versus HGSC. (F) Hallmark gene set enrichment analysis for the comparison shown in (E). (G) Differential gene expression in the Immunoreactive STIC subtype versus HGSC. (H) Hallmark gene set enrichment analysis for the comparison shown in (G).

We also inferred the molecular mechanisms involved in the transition from a precursor lesion to invasive HGSC by comparing HGSCs to both the proliferative subtype and the proliferative/immunoreactive subtypes, assuming that both subtypes were immediate precancerous lesions to invasive carcinomas (Fig. 2E-2H). We observed that the primary mechanisms related to invasion were attributed to epithelial DEGs involved in the interferon response pathway (*ISG15, MX1, OSA1/2/3, IFIT1/6,* and *BST2*), the extracellular matrix remodeling pathway (*MMP1, MMP11*, *COL1A1, COL4A1*), and the epithelial-mesenchymal transition pathway (*LICAM*, *S100A4*, *YAP1*), along with several cancer-promoting genes including IGF2, among others. A similar set of DEGs was also found by comparing HGSCs to all STIC subtypes (Fig. S3D).

Next, we correlated the PIMD STIC subtypes with patterns of DNA copy number changes. We found that the overall aneuploidy index, defined as the chromosomal sizes showing either DNA copy number gain or loss, was highest in both the proliferative and immunoreactive subtypes. Their index levels were comparable to those observed in HGSCs. In contrast, dormant STICs displayed the lowest aneuploidy index, with the mixed subtypes falling in between the two (Fig. 3A). Alongside the extent of aneuploidy, we found that PIMD STIC subtypes differed in the aneuploidy patterns across various chromosomal regions (Fig. 3B). Among the genome-wide copy number changes, the proliferative and dormant subtypes showed the most and least frequent gains or amplifications of the genomic foci harboring *CCNE1* and *MYC,* which are known to characterize HGSCs (Table 1). Fig. 3B demonstrated DNA copy number gain and loss in various chromosomal loci that harbored known cancer driver genes.

**Figure 3.**
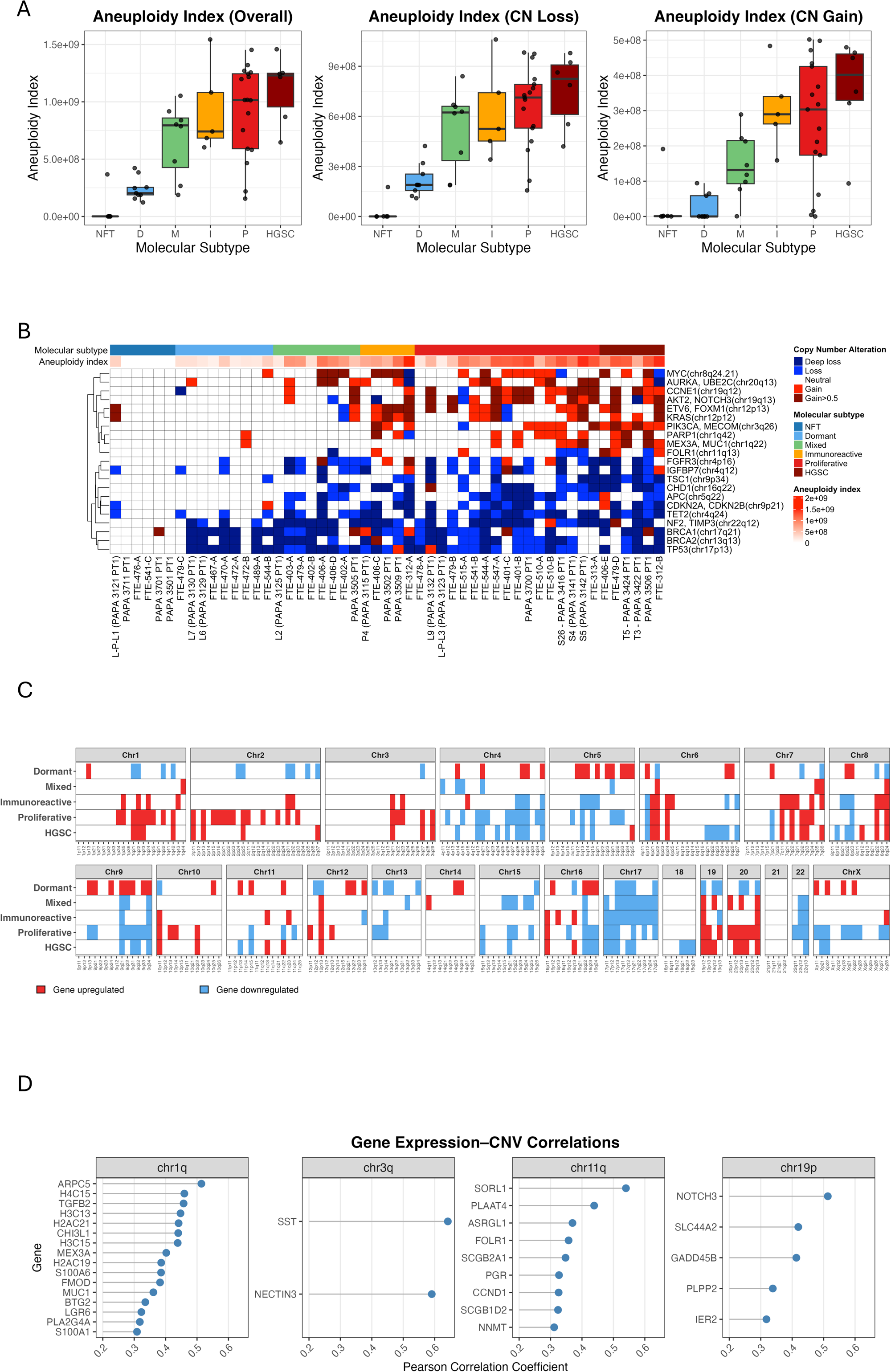
Aneuploidy patterns across molecular subtypes of STICs. (A) The aneuploidy index across all subtypes. (B) The heatmap displays DNA copy number alterations in different STIC subtypes. Representative cancer-associated genes located in the loci exhibiting copy number gain or loss are indicated. (C) Correlation between DNA copy number gain or loss and the up- or down-regulation of mRNA corresponding to genes situated in their cytobands. (D) A list of genes whose copy number alterations significantly correlate with their mRNA expression levels. CN: copy number; CNV: copy number variations; D: dormant; M: mixed; I: immunoreactive; P: proliferative; HGSC: high-grade serous carcinoma.

**Table 1.** Clinicopathological features of precancerous lesions.

|  |  | **Dormant** | **Mixed** | **Immunoreactive** | **Proliferative** | **p-value** |
| --- | --- | --- | --- | --- | --- | --- |
| **No. Patient** |  | 41 | 52 | 26 | 47 |  |
| **Age at the time of diagnosis** |  |  |  |  |  | <0.001 |
|  | 30-39 | 0 (0.0) | 0 (0.0) | 1 (3.8) | 0 (0.0) |  |
|  | 40-49 | 3 (7.3) | 0 (0.0) | 5 (19.2) | 0 (0.0) |  |
|  | 50-59 | 28 (68.3) | 15 (28.8) | 9 (34.6) | 9 (21.4) |  |
|  | 60-69 | 9 (22.0) | 9 (17.3) | 4 (15.4) | 20 (47.6) |  |
|  | 70-79 | 1 (2.4) | 28 (53.8) | 7 (26.9) | 13 (31.0) |  |
| **BRCA germline mutation status** |  |  |  |  |  | 0.001 |
|  | Negative | 30 (73.2) | 36 (69.2) | 12 (46.2) | 41 (87.2) |  |
|  | BRCA1 | 0 (0.0) | 6 (11.5) | 8 (30.8) | 2 (4.3) |  |
|  | BRCA2 | 6 (14.6) | 6 (11.5) | 5 (19.2) | 2 (4.3) |  |
|  | Other | 5 (12.2) | 4 (7.7) | 1 (3.8) | 2 (4.3) |  |
| **Nuclear atypia** |  |  |  |  |  | <0.001 |
|  | Low-grade | 32 (78.0) | 24 (46.2) | 4 (15.4) | 4 (8.5) |  |
|  | High-grade | 9 (22.0) | 28 (53.8) | 22 (84.6) | 43 (91.5) |  |
| **Architectural subtype (surface)** |  |  |  |  |  | <0.001 |
|  | Flat | 39 (95.1) | 39 (75.0) | 19 (73.1) | 26 (55.3) |  |
|  | BLAD | 2 (4.9) | 13 (25.0) | 7 (26.9) | 21 (44.7) |  |
| **Stromal lymphocytes** |  |  |  |  |  | 0.031 |
|  | Non-enriched | 39 (95.1) | 42 (80.8) | 19 (73.1) | 43 (91.5) |  |
|  | Enriched | 2 (4.9) | 10 (19.2) | 7 (26.9) | 4 (8.5) |  |
| **Ki67 index** | Low (<5%) | 13 (32.5) | 19 (36.5) | 5 (19.2) | 3 (6.4) | <0.001 |
|  | Intermediate (5-20%) | 22 (55.0) | 16 (30.8) | 8 (30.8) | 15 (31.9) |  |
|  | High (>20%) | 5 (12.5) | 17 (32.7) | 13 (50.0) | 29 (61.7) |  |
| **p53 IHC pattern** | Missense | 32 (78.0) | 44 (84.6) | 24 (92.3) | 35 (74.5) | 0.522 |
|  | Loss/null | 8 (19.5) | 8 (15.4) | 2 (7.7) | 11 (23.4) |  |
|  | Wildtype | 1 (2.4) | 0 (0.0) | 0 (0.0) | 1 (2.1) |  |
| **CCNE1 copy number gain** |  | 0/9 (0.0) | 2/8 (25.0) | 1/5 (20.0) | 10/17 (58.8) | 0.017 |
| **MYC copy number gain** |  | 1/9 (11.1) | 3/8 (37.5) | 3/5 (60.0) | 7/17 (41.2) | 0.275 |

IHC: immunohistochemistry; BLAD: budding, loosely attached, and detached STIC cells

### Gene Expression Mapping to Chromosomes

Among the samples with available transcriptomic and aneuploidy data, we observed statistically significant co-upregulation of genes located in subchromosomal regions (cytobands as the “zip code”), showing DNA copy number gains and co-loss in regions demonstrating DNA copy number loss (Fig. 3C). Our findings highlighted consistent downregulation of genes on chromosomes 17 and 22 in STIC lesions exhibiting chromosomal deletions. Additionally, most dormant STICs co-downregulated genes located on chromosome 13, which were often lost. Conversely, upregulation of 8q24 containing *MYC* was evident in proliferative, immunoreactive, mixed STICs, and HGSC, supporting its role in tumor progression. We further demonstrated that the STIC-associated genes had a significant correlation between their RNA levels and DNA copy numbers (Fig. 3D), indicating that DNA copy number gain likely contributed to the increased gene expression observed in many STIC specimens.

### Clinicopathological Features of the STIC Subtypes

Since STICs were discovered and originally defined by histopathologic examination, we determined how the PIMD molecular subtypes correspond to their pathological features. We focused on the severity of cytological atypia and architectural features, both of which serve as morphological hallmarks of precancerous and malignant transformation recognized in routine pathology. High-grade STICs exhibited prominent nuclear atypia, enlargement, and stratification, along with occasional mitotic figures or apoptotic bodies. In contrast, the observed severity of these features was much less in low-grade STICs. Some pathologists classified these low-grade lesions as serous tubal intraepithelial lesions (STIL). Based on this binary grading system, we found that the percentage of high-grade STICs was highest in the proliferative subtype (91.5%), followed by the immunoreactive subtype (84.6%), the mixed subtype (57.1%), and lowest in the dormant subtype (22.5%) (Table 1). The absence of high-grade nuclear features, combined with transcriptional similarity to normal epithelium (Fig. 1B), suggested that the dormant subtype is evolutionarily distinct from other STICs. Therefore, we performed clinicopathological correlation in the proliferative, immunoreactive, and mixed STIC subtypes (Table 1).

Most notably, pathogenic germline *BRCA1/2* mutations (g-*BRCA1/2^mut^*) were enriched in immunoreactive subtypes; g-*BRCA1/2^mut^*were present in half of immunoreactive STICs. The other molecular subtypes showed a significantly lower frequency of such mutations (Fig. 5A). With this finding, we compared the transcriptomes of STICs derived from women carrying g-*BRCA1/2^mut^* with those from g-*BRCA1/2^wildtype^*STICs (Fig. 4A to 4C). The findings provided validation, as the observed patterns paralleled those seen in comparisons of immunoreactive STICs with other subtypes. Several immunoregulatory genes and antigen-presenting pathways were found to be upregulated in g-*BRCA1/2^mut^* STICs and the immunoreactive subtype (Fig. 4C). However, these changes were absent from their corresponding histologically unremarkable fallopian tube epithelium.

**Figure 4.**
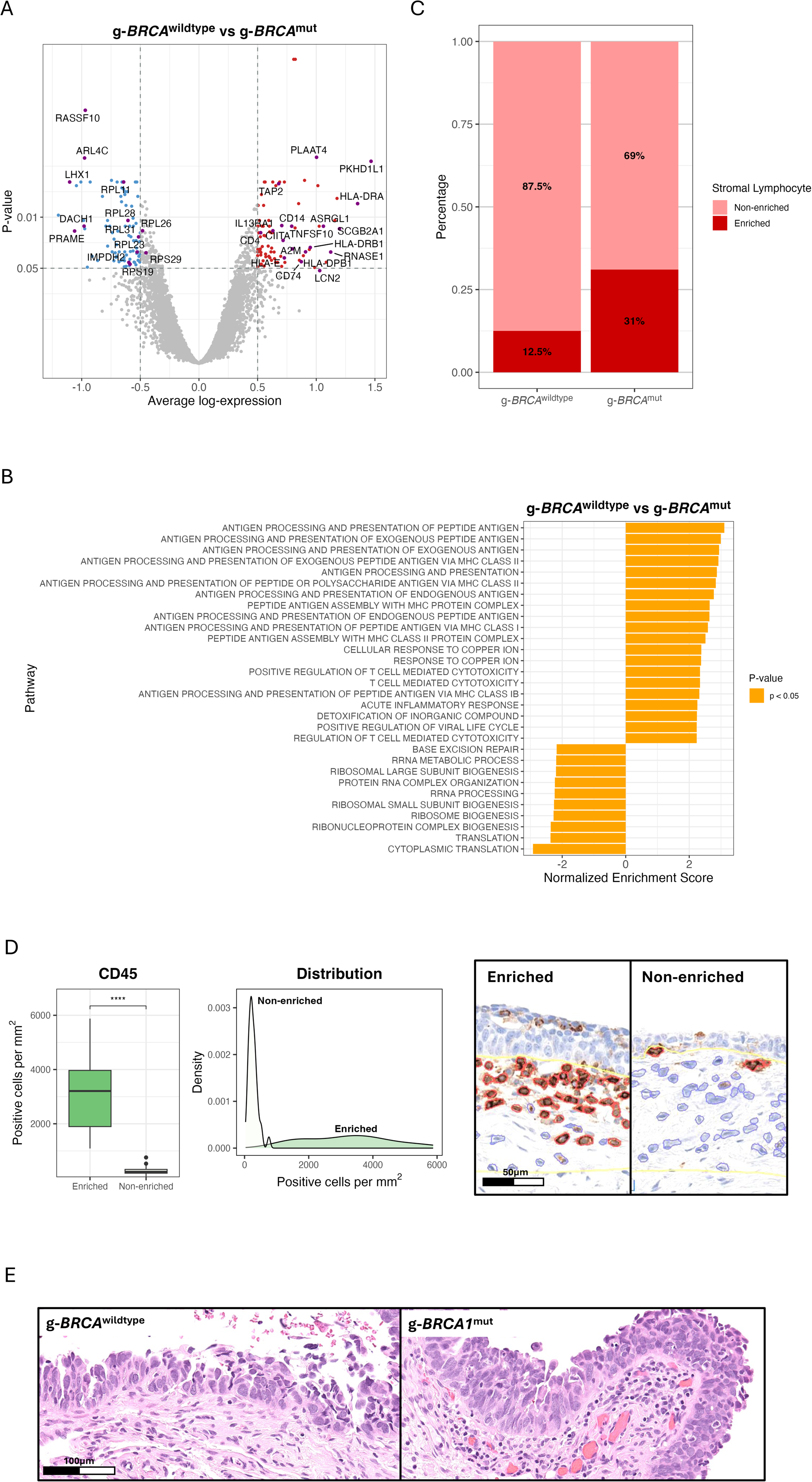
Enrichment of immune cells in the STICs harboring *BRCA1/2* germline mutations. (A) Volcano plot showing differentially expressed genes in germline (g)-*BRCA1/2^mut^* mutation STICs, compared to the g-*BRCA1/2^wildtype^*STICs. (B) Pathway analysis based on the differentially expressed genes from (A). (C) Percentages of STICs show enriched stromal lymphocytes. (D) CD45L cell density per mm² in pathologist-defined lymphocyte-enriched versus non-enriched stroma (left); smoothed density plot displaying the distribution of CD45⁺ cell densities across all regions (middle); representative CD45^+^ cells with automated segmentation overlays from an enriched region and a non-enriched region (right panels). (E) Representative STIC lesions illustrate stromal lymphocyte density: non-enriched in a g-*BRCA1/2^wildtype^*sample (left) versus enriched infiltrate in a g-*BRCA1^mut^* sample (right).

**Figure 5.**
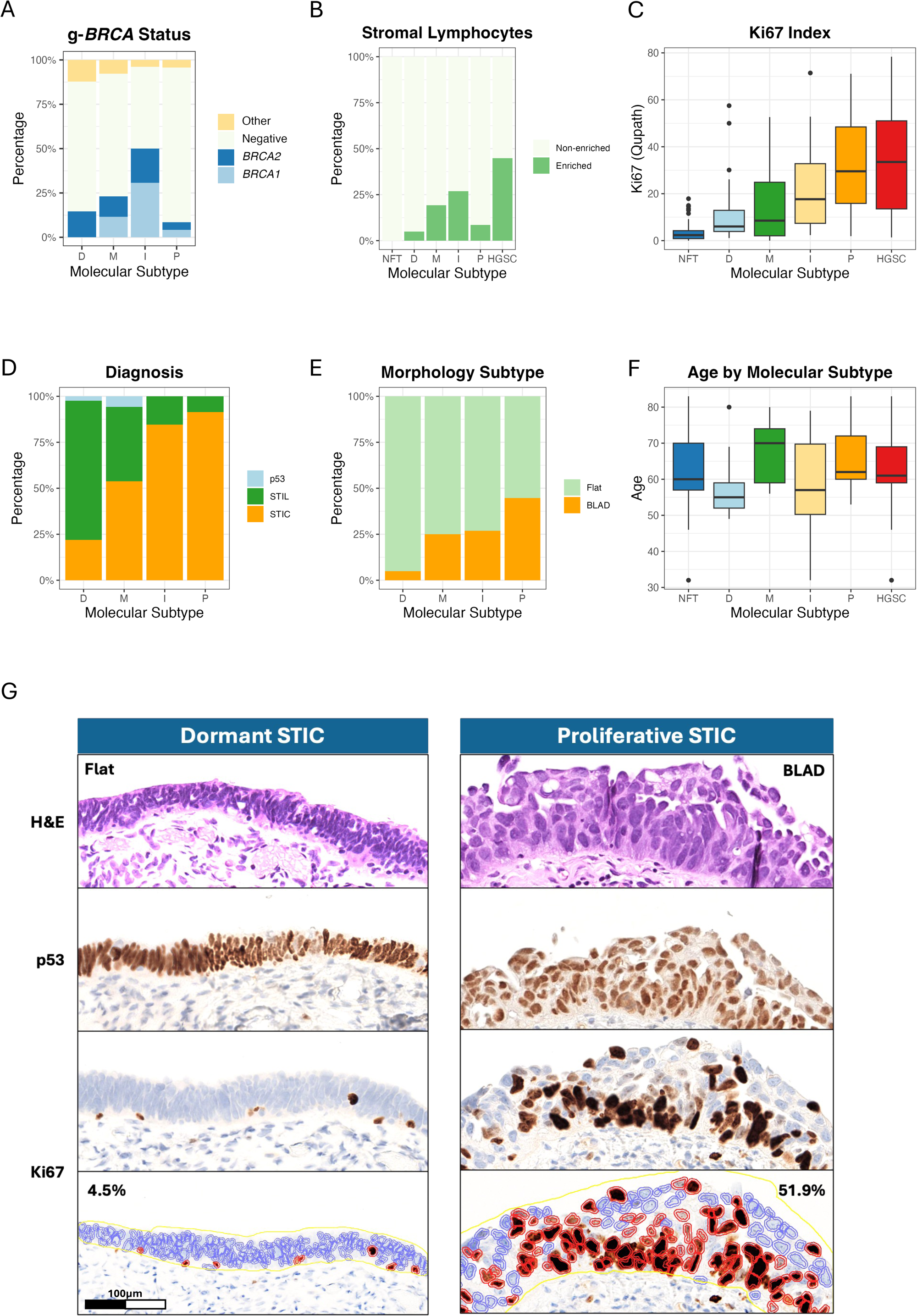
Correlation of STIC molecular subtypes with clinical and pathological features. Correlations with (A) germline*-BRCA1/2* mutation status, (B) stromal lymphocyte infiltration, (C) proliferative activity based on the Ki67 labeling index, (D) pathology diagnosis, (E) morphological features, and (F) age at presentation. (G) Photomicrographs of H&E-stained, p53-stained, and Ki67-stained slides contain one STIC of the dormant subtype and another of the proliferative subtype. Compared to the dormant STIC, the proliferative STIC shows budding, loosely adherent, and detached (BLAD) tumor cells and has a Ki67 labeling index of 51.9% (4.5% in dormant STIC), quantified by the QuPath program (bottom panels). P: proliferative subtype; I: immunoreactive subtype; M: mixed subtype; D: dormant subtype.

Microscopic evaluation demonstrated that, when compared to STICs, HGSCs had more lesions with prominent lymphocytic infiltrates in the stroma (Fig. 5B). Based on this observation we then investigated whether there was variability in enriched lymphocytes across the molecular subtypes of STICs compared to corresponding normal fallopian tube mucosa. We applied the diagnostic pathology criteria used in clinical care to define lymphocyte enrichment. Our morphology-based classification was validated on a subset of CD45-immunostained slides. As shown in Fig. 4D, pathologist-identified lymphocyte-enriched regions exhibited a significantly higher density of CD45-positive cells (p<0.001). We found enriched lymphocytic infiltrate (compared to the adjacent normal fallopian tube) in 61.5% of immunoreactive STICs. We also independently observed lymphocytic enrichment in g-*BRCA1/2^mut^* STICs. Examples of stromal immune cells from g-*BRCA1/2^mut^* STICs and g-*BRCA1/2^wldtype^* STICs are shown in Fig. 4E. Importantly, the number of cases with lymphocytic enrichment in histologically unremarkable tubal epithelium from either g-*BRCA1/2^mut^* STICs or the g-*BRCA1/2^wildtype^* STICs did not significantly differ.

Proliferative activity, measured by the Ki67 labeling index (% of positive epithelial cells), was assessed by both manual counting and QuPath software, which were highly correlated (r^2^= 0.94). We found the index to be highest in the proliferative subtype, followed by immunoreactive, mixed, and dormant STICs in decreasing order (Fig. 5C and 5G).

We also observed that both proliferative and immunoreactive subtypes corresponded to pathologically defined “STICs” (Fig. 5D). Evaluation of their architectural features was based on a recent morphology-based study that reported two architecturally distinct groups of STICs ^18^. One group exhibited a flat and cohesive surface (Flat) of the STIC lesion, while the other was characterized by STIC cells displaying a budding, loosely adherent, or detached (BLAD) morphology (examples shown in Fig. S4A), suggestive of a potentially increased risk of dissemination. The BLAD feature was observed in 21 (44.7%) of 47 proliferative STICs and in only 2 (4.9%) of 41 dormant STICs (Fig. 5E). Both immunoreactive and mixed subtypes exhibited BLAD STIC percentages between these two extremes. In a separate analysis, independent of PIMD subtypes, BLAD STICs had a higher Ki67 index. BLAD STICs were also characterized by increased stromal lymphocyte infiltration, along with elevated expression of *SOX4*, *FUT8*, *APOA1*, and *TNNT1* compared to Flat STICs (Fig. S4B to S4D). Hallmark pathway analysis revealed that BLAD STICs, compared to their Flat counterparts, exhibited increased activity in E2F targets, G2M checkpoint, mitotic spindle, MYC targets, DNA repair, and epithelial-mixed transition (EMT) pathways (Fig. S4E). In general, patients’ ages at the time of diagnosis were similar across the different subtypes, despite the slightly older age at diagnosis with mixed subtype (Fig. 5F).

### Validation of Representative Markers

We selected four representative genes: *SOX4*, *MCM7*, *DBN1*, and *KIFC1,* which were upregulated in STICs and HGSCs according to spatial transcriptomic analysis, for orthogonal validation at the tissue level. These genes were chosen based on the following criteria: 1) upregulation in both STICs and HGSCs, but not in normal fallopian tube epithelium or stroma; 2) potential biological significance in tumor development; 3) availability of antibodies suitable for formalin-fixed and paraffin-embedded tissue sections; and 4) genes previously unreported in STIC lesions. We performed immunohistochemistry to validate their expression patterns in a cohort that included many lesions not found in the original discovery set. Consequently, we compared the protein expression levels among various STIC subtypes. Fig. 6A presents photomicrographs of immunostaining results obtained with each of the four proteins. *DBN1* protein expression was membranous, with positive signals detected in almost all STIC and HGSC cells in both specimens. Immunoreactivity for *MCM7*, *SOX4*, and *KIFC1* was observed solely in the nuclei of STIC and HGSC cells, as expected.

**Figure 6.**
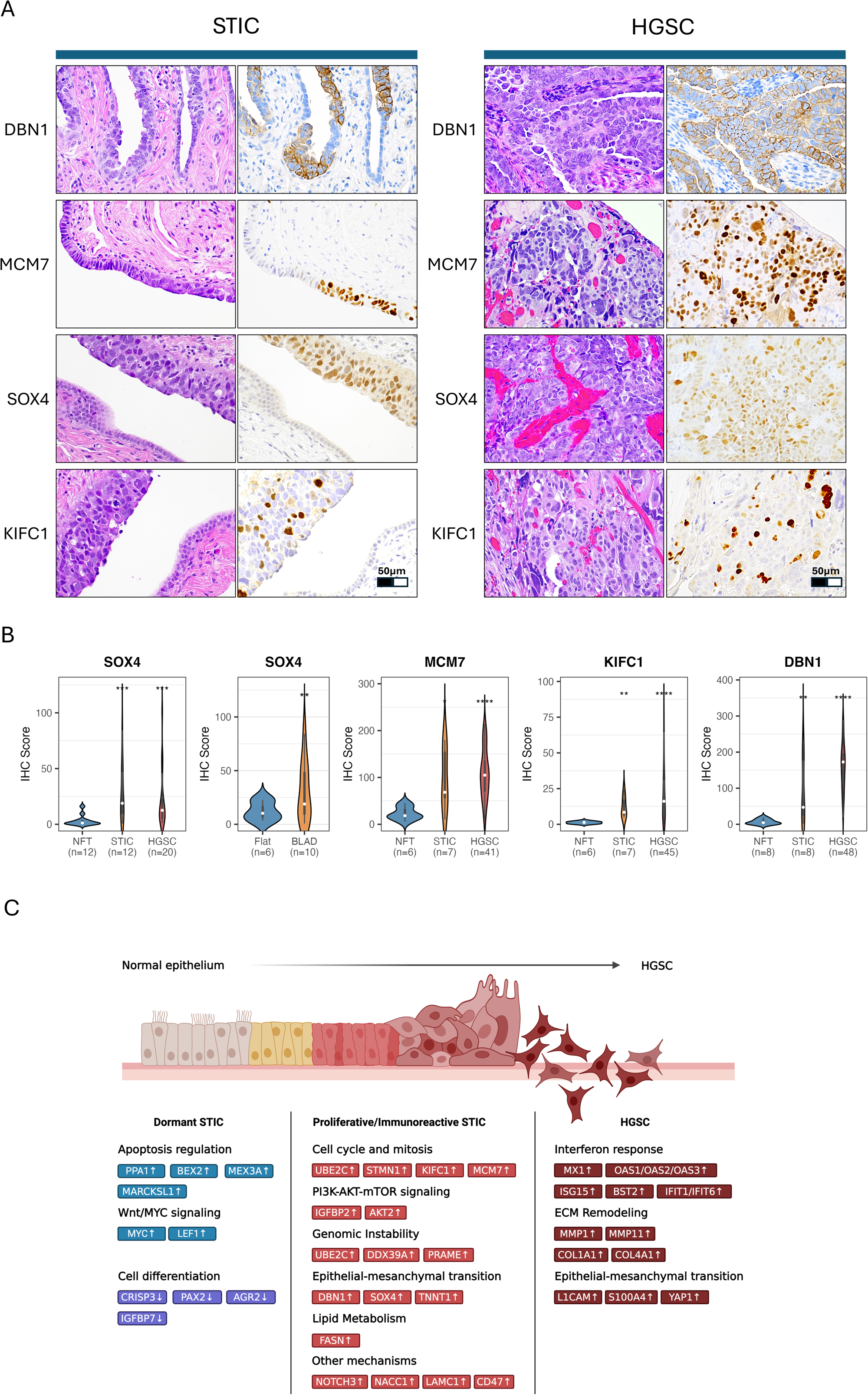
Validation of upregulated genes in STICs and HGSCs. (A) H&E-stained and immunostained images for SOX4, MCM7, KIFC1, and DBN1 in selected STIC and HGSC samples. (B) Quantification of protein expression using the QuPath program. Violin plots display IHC scores for the selected markers across different diagnostic groups and morphologic categories. (C) Schematic illustration of the progression from normal fallopian tube epithelium (NFT) through serous tubal intraepithelial carcinoma (STIC) to high-grade serous carcinoma (HGSC). The possible genes and the pathways in which they are involved are listed.

Digital pathology quantified protein expression levels using QuPath and found that the immunohistochemistry scores of *SOX4*, *MCM7*, *KIFC1*, and *DBN1* in STICs and HGSCs were significantly higher than those in normal fallopian tube epithelium (Fig. 6B). Furthermore, *SOX4* expression levels were significantly elevated in STICs exhibiting the BLAD morphology, which is a morphology more commonly observed in the proliferative, immunoreactive, and mixed subtypes than in STICs with flat surface morphology. In this cohort, we did not find significant differences in the ages when the specimens were obtained.

## Discussion

Prevention and early detection hold the keys to reducing cancer morbidity and mortality. The updated paradigm regarding the origin of high-grade serous carcinoma (HGSC) is transforming multiple facets of ovarian cancer research and clinical practice. Most notably, opportunistic salpingectomy, a procedure aimed at reducing ovarian cancer risk by excising fallopian tubes during another elective intra-abdominal surgery, has gained increasing attention as an efficacious and potential primary prevention method for HGSC in the general population (14,22-24). Second, early salpingectomy followed by delayed oophorectomy closer to the natural age of menopause is being investigated as an alternative to traditional salpingo-oophorectomy starting as early as age 35, in order to delay loss of ovarian endocrine function in women at increased risk of developing HGSC, including those with inherited pathogenic mutations in *BRCA1, BRCA2,* and other homologous recombination genes. Third, studies utilizing liquid Pap cervical samples, uterine lavage, and innovative imaging modalities promise early detection of STICs and incipient HGSC in the fallopian tubes (25-27). Identifying tubal pre-cancerous-specific markers applicable to these early detection and imaging approaches would open new avenues and propel the advancement of this research area.

In this study, we report four distinct molecular classes of HGSC precursors: Proliferative, Immunoreactive, Mixed, and Dormant STIC subtypes (the PIMD classification), that are characterized by unique molecular and clinicopathological features. Our data presented herein not only shed new light on the earliest molecular changes in ovarian cancer but also have several translational implications.

First, during the progression of HGSC, various molecular mechanisms evolve as early as the precursor stages, indicating multiple initiating events for HGSC. Among these, STICs that originated from the fallopian tubes of germline *BRCA1/2* pathogenic mutation (g-*BRCA^mut^*) carriers are unique because they are more likely to elicit an immune-related phenotype, including upregulation of antigen-presenting genes, activation of interferon pathways, and an increase in stromal lymphocytes. Interestingly, no significant difference in gene expression changes occurred in normal fallopian tube epithelium between g-*BRCA^mut^* and g-*BRCA^wildtype^* specimens. These results suggest that the molecular genetic alterations involved in STIC transformation are likely necessary to trigger immune activation.

Indeed, molecular genetic data reveal one possible mechanism. Almost all STICs, including those containing *g-BRCA1/2^mut^*, not only harbored *TP53* mutations but also lost one chromosome 17 and one chromosome 13, based on our RealSeqS analysis. Since both *TP53* and *BRCA1* reside on chromosome 17, the deletion of this chromosome ostensibly leads to biallelic inactivation of both genes, thereby creating a double “two-hit” scenario predisposing to carcinogenesis. In contrast, *g-BRCA^wildtype^* STICs may retain an active *BRCA1* allele that continues to maintain BRCA1-mediated homologous recombination DNA repair to a certain degree. The loss of chromosome 13 is also noteworthy because it harbors *BRCA2*. Consequently, STICs from g-*BRCA2 ^mut^* may likely manifest biallelic inactivation, resulting in homologous recombination deficiency.

Second, our lack of knowledge about STIC biology and outcomes has prevented development of clinical guidelines for managing patients with incidental STIC diagnoses. Whether women diagnosed with a STIC require exploratory laparoscopy, liquid biopsy, chemotherapy, mutant p53 reactivator ^40^, or simply close clinical follow-up (and how to do that effectively) remains unclear. A meta-analysis reported a gradual increase in the incidence of disseminated HGSC from incidental STICs, with an incidence of HGSC reaching 27.5% after 10 years of follow-up ^41^. In those unfortunate cases, it can be envisioned that before salpingectomy, detached STIC cells from the fallopian tube surface may have disseminated to peritoneal soft tissues, where they subsequently establish peritoneal carcinomatoses. The results of this report support the heterogeneity of STICs and underscore the need to develop a risk stratification strategy. This task has become increasingly urgent due to the anticipated rise in STIC diagnoses following national and international advocacy for opportunistic salpingectomy as a primary prevention strategy for HGSC^42^. Our work reveals that both proliferative and immunoreactive STIC subtypes display several features indicative of increased aggressiveness. Specifically, these STICs are more likely to show pronounced chromosomal instability, amplify *CCNE1* and *MYC*, overexpress multiple cancer-promoting genes, show increased levels of proliferative activity, exhibit high-grade dysplasia, and have a BLAD morphology suggestive of cell shedding from STIC surfaces.

Third, dormant STICs represent a biologically intriguing subtype because they are clonally developed and harbor *TP53* mutations, yet they appear to lack the features seen in proliferative and immunoreactive subtypes. Most do not proliferate as much as their normal fallopian tube epithelial cells. Among all PIMD subtypes, dormant STICs are molecularly closest to normal fallopian tube epithelium and furthest from HGSC. They exhibit less aneuploidy than other STIC subtypes and express genes associated with tubal epithelium differentiation. Despite similarities to other subtypes that lose chromosomes 13, 17, and 22, dormant STICs may have reached an evolutionary impasse and may no longer proliferate to create more genetic or transcriptomic variants for selection. This finding suggests that the loss of these chromosomes is necessary but insufficient to drive tumor progression. Severe telomere attrition and/or oncogene-induced activation of senescent programs has been reported in some STICs compared to HGSCs ^43,44^. In one study, all incidental STICs with low proliferative activity (Ki67 <5%) show severe telomere shortening ^43^. Alternatively, dormant STICs may not be completely dormant; They may represent a temporarily quiescent state awaiting reactivation through increased transposon activity or alterations in their microenvironment ^1^. Whether dormant STICs are clinically indolent remains to be studied in future STIC consortia that will gather sufficient cases and long-term follow-up information.

Fourth, we identified possible mechanisms that drive STIC lesions to become invasive. The aneuploidy index, as indicated by genome-wide DNA copy number alterations, does not significantly differ between the proliferative and/or immunoreactive subtypes STIC and HGSC. Gene expression related to cell cycle control does not vary either. However, the main difference lies in the upregulation of genes associated with the interferon response pathway, the epithelial-mesenchymal transition pathway, and the extracellular matrix remodeling pathway (Fig. 6C). Moreover, in addition to epithelial cells, the HGSC stroma also activates the interferon response pathway and enriches tumor-associated macrophages that modulate tumor immunity ^45^. These findings suggest that invasion likely occurs through transcriptomic reprogramming to promote crosstalk between STIC and its microenvironment.

Fifth, the mixed STIC subtypes included STICs showing either low-grade or high-grade nuclear atypia, along with low or high proliferative activity. The overall aneuploidy index of the mixed type lies between dormant and proliferative/immunoreactive subtypes. Chromosomal deletions are more pronounced than those in dormant STICs, while chromosomal gains are less frequent than in proliferative/immunoreactive subtypes. It is possible that additional DNA copy number gains, particularly in loci containing tumor-promoting genes, are crucial for progression to proliferative/immunoreactive STICs, the presumably immediate precursors of HGSC. Conversely, “mixed” subtypes may signify a biologically distinct group of lesions beyond their denotation.

Lastly, since morphology-based diagnosis is the standard of patient care, our data suggest a binary pathology grading system: low-grade STIC versus high-grade STIC. Low-grade STICs mainly belong to the dormant subtype, whereas high-grade STICs encompass the majority of proliferative and immunoreactive subtypes. This proposition is analogous to the Bethesda grading system used in clinical pathology for evaluating cervical lesions ^46^. This system classifies cervical precancerous lesions into low-grade and high-grade squamous intraepithelial lesions and has been used for decades to guide clinical decisions. Moreover, based on an integrated analysis of STIC lesions, we propose a companion marker panel that can assist in the differential diagnosis of challenging cases to determine whether the minute area of concern is an STIC and, if so, its grade. In addition to Ki67 and p53 immunohistochemistry, we suggest immunohistochemistry using antibodies against the proteins encoded by *MCM7*, *DBN1*, and *KIFC1* in combination. Their mRNA levels are significantly associated with DNA copy number gain, indicating a selective advantage in tumor progression. Indeed, the proteins encoded by *MCM7* ^47^, *DBN1* ^48^, and *KIFC1* ^49^ are known to participate in tumor development. Importantly, these markers have been validated in this study using immunohistochemistry, and the antibodies are commercially available.

While the current study offers exciting new insights into the earliest molecular changes in HGSC initiation, it is important to acknowledge its limitations. Foremost, longer follow-up data (greater than 4 years) are largely lacking in our current cohort. Nevertheless, the PIMD classification provides a foundation for future collaborative studies on clinical correlation. This is indeed an ongoing area of intense investigation for us as actionable risk stratification requires the identification of relevant clinical correlates. Second, unlike carcinomas or precursor lesions of other tissue types, STIC lesions are generally not larger than 1 mm and are detected microscopically only in formalin-fixed and paraffin-embedded tissue sections. Because of their small size, many of the lesions are exhausted at deeper tissue levels, making it not always feasible to perform multi-omics on the same lesions. Fortunately, we can achieve this task in many cases due to the relatively large number of STICs collected.

In summary, integrated spatial analysis of this study sheds new light on the molecular landscape of transcriptomics and aneuploidy patterns alongside several clinical and pathological characteristics in ovarian precursor lesions (Fig. 6C). Our integrated and spatial analysis reveals that there are various subtypes of precursor lesions that fall under the rubric of STIC, each exhibiting unique biological features. Our data will help create a publicly accessible database that can be used to investigate specific genes and conduct novel computational analyses to explore ovarian serous carcinogenesis.

## Acknowledgements

This study is supported by the Richard W. TeLinde Endowment from the Johns Hopkins University, NIH/NCI (U2CCA271891, P50CA228991, R01CA260628, and R01CA215483), and Tina Wish Foundation. The Nanostring DSP GeoMx study was supported by the Break Through Cancer (BTC-IOC).

