## Supplemental Figure for "Integrated Spatial Analysis of Ovarian Precancerous Lesions"

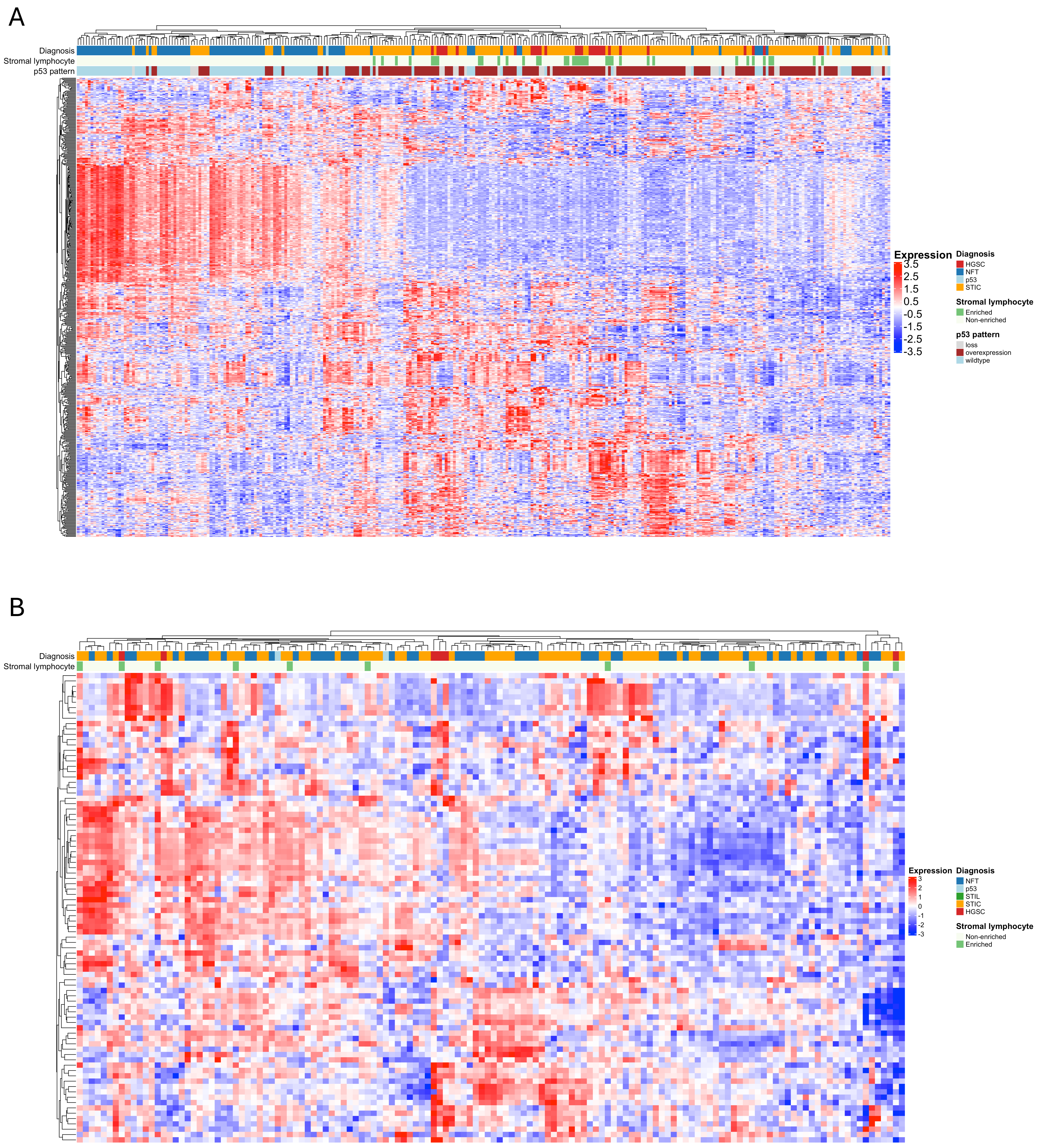


**Supplemental Figure 1. Heatmap.** (A) Unsupervised heatmap with hierarchical clustering of epithelial samples based on highly variable genes (variance > 0.9). (B) Unsupervised heatmap with hierarchical clustering of stromal samples based on highly variable genes (variance > 0.9).

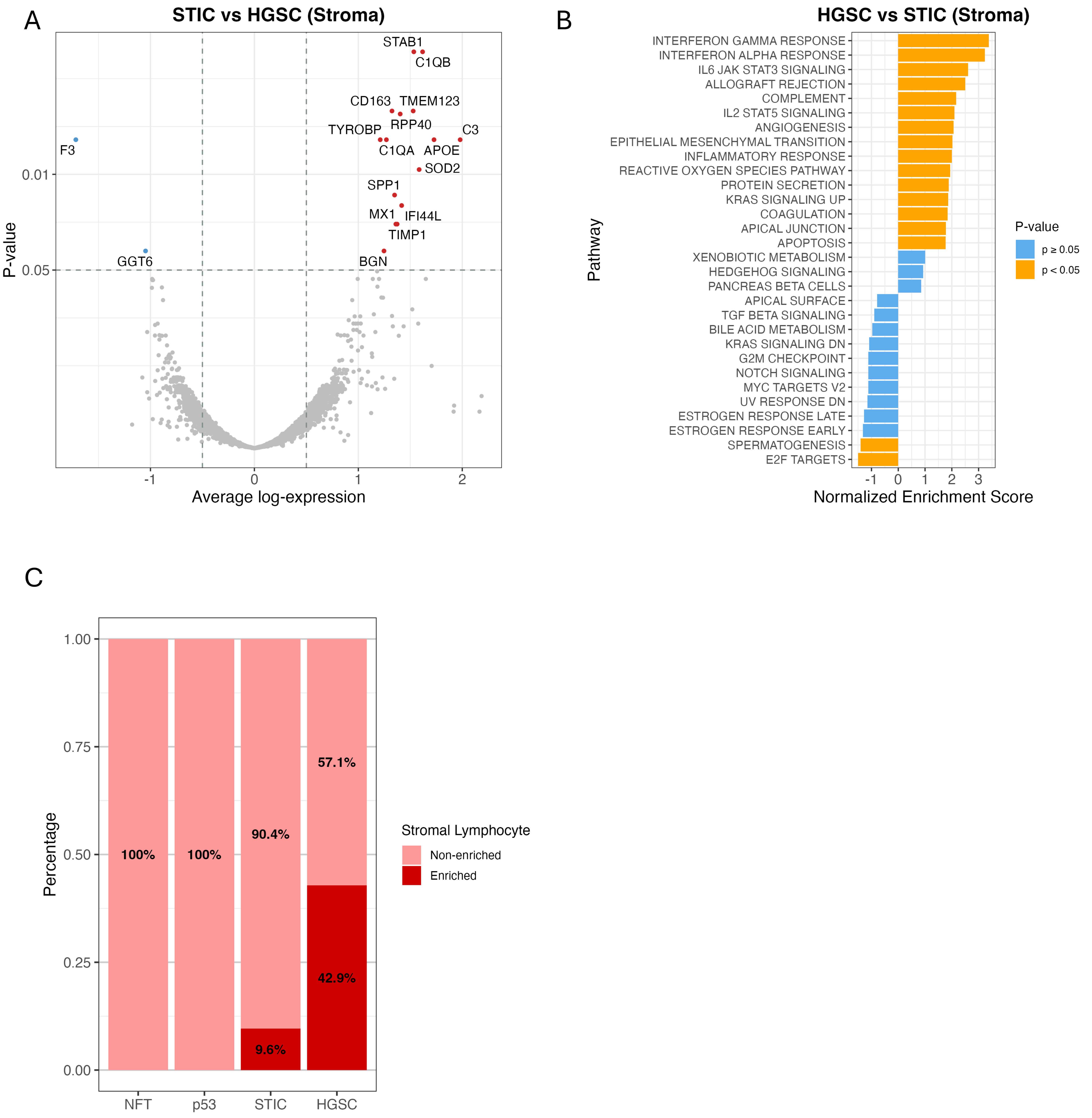


**Supplemental Figure 2. Differential expression analysis of stromal components.** (A) Volcano plot showing differentially expressed genes between stromal samples of STIC and HGSC. (B) Pathway analysis identifying enriched pathways in HGSC vs. STIC stroma. (C) Distribution of stromal lymphocytes across different diagnoses compared to background lymphocytes. (D) CD45+ immune cells in an STIC show enriched lymphocytes compared to the area of the adjacent (background) normal fallopian tube.

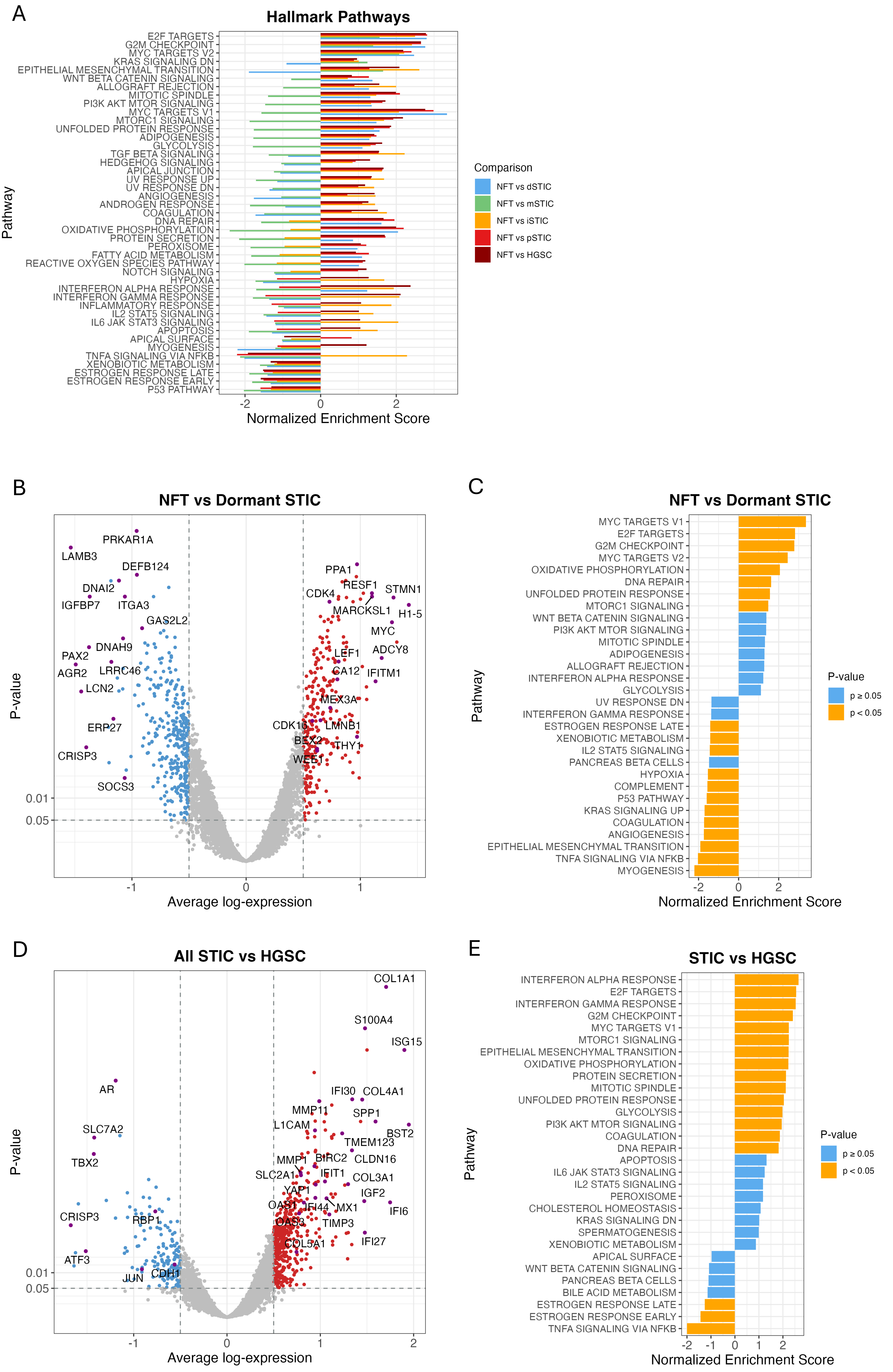


**Supplemental Figure 3. Differential expression analyses.** (A) Hallmark gene-set enrichment across all STIC molecular subtypes versus normal fallopian tube epithelium (NFT). (B) Differential genes expression in the Dormant STIC subtype versus NFT. (C) Hallmark gene-set enrichment analysis for the Dormant STIC versus NFT comparison shown in (B). (D) Differential genes expression in pooled STIC samples (all subtypes) versus HGSC. (E) Hallmark gene-set enrichment analysis for the pooled STIC versus HGSC comparison shown in (D).

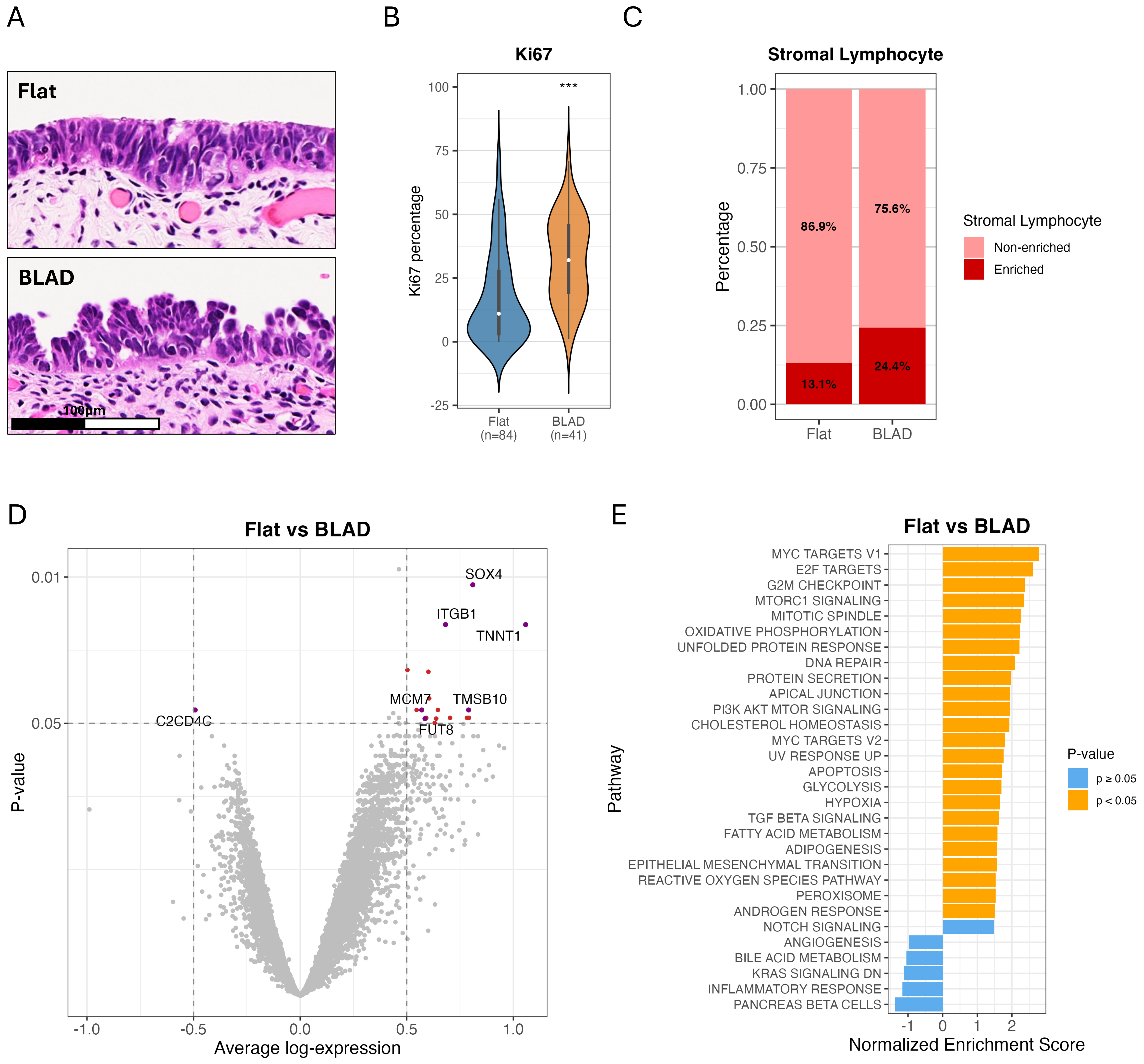


**Supplemental Figure 4. Transcriptomic and immune cell landscape in STICs showing aggressive and indolent morphologies.** (A) Representative H&E images of a BLAD STIC (bottom) and a Flat STIC (top). (B) Boxplot illustrating Ki67 expression labeling percentage in BLAD and Flat lesions. *** $p < 0.005$. (C) Percentages of STICs enriching tumor-infiltrating lymphocytes (TIL) in BLAD and Flat lesions. (D) Volcano plot comparing differentially expressed genes between BLAD and Flat morphologies. Significant genes are indicated. (E) Hallmark pathway analysis highlights key pathways differentially regulated between BLAD and Flat lesions.
